# Fast Local Alignment of Protein Pockets (FLAPP): A system-compiled program for large-scale binding site alignment

**DOI:** 10.1101/2022.07.13.499925

**Authors:** Santhosh Sankar, Naren Chandran Sakthivel, Nagasuma Chandra

## Abstract

Protein function is a direct consequence of its sequence, structure and the arrangement at the binding site. Bioinformatics using sequence analysis is typically used to gain a first insight into protein function. Protein structures, on the other hand, provide a higher resolution platform into understanding functions. As the protein structural information is increasingly becoming available through experimental structure determination and through advances in computational methods for structure prediction, the opportunity to utilize this data is also increasing. Structural analysis of small molecule ligand binding sites in particular provide a direct and more accurate window to infer protein function. However it remains a poorly utilized resource due to the huge computational cost of existing methods that make large scale structural comparisons of binding sites prohibitive. Here we present an algorithm called FLAPP that produces very rapid atomic level alignments. By combining clique matching in graphs and the power of modern CPU architectures, FLAPP aligns a typical pair of binding site binding sites at ~12.5 milliseconds using a single CPU core, ~ 1 millisecond using 12 cores on a standard desktop machine, and performs a PDB-wide scan in 1-2 minutes. We perform rigorous validation of the algorithm at multiple levels of complexity and show that FLAPP provides accurate alignments. We also present a case study involving vitamin B12 binding sites to showcase the usefulness of FLAPP for performing an exhaustive alignment based PDB-wide scan. We expect this tool will be invaluable to the scientific community to quickly align millions of site pairs on a normal desktop machine to gain insights into protein function and drug discovery for drug target and off-target identification, and polypharmacology.

## Introduction

Proteins are the primary architects of biological processes and study of their three-dimensional structures provide a means to understand them at the highest resolution. Protein function has traditionally been elucidated by biochemical assays, but the genomic era has provided us with protein sequences at an unprecedented rate, leading to a large gap between protein identification and understanding their function. Computational approaches allow us to bridge the gap. Typically, such computational approaches utilize various measures of similarity inferred to transfer knowledge from annotated proteins^1^. Sequence-based comparison methods such as BLAST offer a great heuristic in rapidly identifying sets of proteins that perform similar functions^2^. On the other hand, structural alignment evaluates similarity by considering their three-dimensional shapes. While alignments that use fold information are more effective than traditional sequence alignments, those that assess similarity in the binding sites provide a higher resolution and more direct means to function assignment.

Many algorithms have been reported for comparing binding sites. These broadly fall into: i) alignment-free and ii) alignment-based. Alignment-free comparison methods use physicochemical descriptors (eg.hydrogen bonding patterns) for assessing similarity between binding sites^3–5^, where-as alignment-based methods aim to generate 3D alignments that give residue-residue correspondences between the sites (eg., PocketAlign, G-LoSA, SiteMotif and SiteEngine^6–9^). Due to the sequentially discontinuous nature of the binding sites, popular sequence or fold alignment algorithms such as dynamic programming are not readily amenable for aligning binding sites. Hence, alignment-free methods are usually preferred for this purpose. Despite the speed of alignment-free methods (eg., PocketMatch, RAPMAD), alignment-based methods (eg.,PocketAlign) are preferred because they provide residue-level alignments. However, their runtime performance remains a bottleneck for their application in large-scale site comparison exercises. For example, SiteEngine (2005) and PocketAlign (2011), take ~2 minutes to align a pair of sites^6,9^. Alternatives such as G-LoSA and SiteMotif are able to bridge this gap by doing one alignment per second or less^7,8^. Nevertheless, they become infeasible when the number of comparisons is in the order of a million or more. This highlights the need for a fast, accurate and scalable site alignment algorithm.

In this work, we present a new algorithm, Fast Local Alignment of Protein Pockets (FLAPP), for rapid and accurate alignment of binding sites, facilitating proteome-wide or PDB-wide scans. The algorithmic design leverages the modern CPU architecture and efficient data structures to generate alignments comparable to state-of-the-art methods. We demonstrate the superiority of FLAPP by benchmarking against the best available methods currently. Lastly, we showcase the efficiency of FLAPP by deriving motifs in Vitamin B12 ligand binding sites and exhaustively scanning the motifs against all binding pockets in PDB.

## Methods

### 2.1 Algorithmic Implementation

A ‘binding site’ is defined as a demonstrated small molecule binding region taken from a protein-ligand complex in the PDB, while a ‘pocket’ refers to a putative small molecule recognition site in a protein. For proteins that are complexed with ligands, the binding site represents those residues that are present within 4.5Å of any ligand atom. In cases where the structure of protein is solved in an apo form (i.e without ligand), then the pockets can be identified using established pocket prediction algorithms such as PocketDepth, FPocket and SiteHound^10–12^. Both ‘pocket’ and the ‘binding site’ represent the similar meaning in the context of this study.

The FLAPP implementation (Fig 1) constitutes three modules: (i) Site Representation ii) Seed selection and (iii) Optimal alignment building.

**Fig 1:**
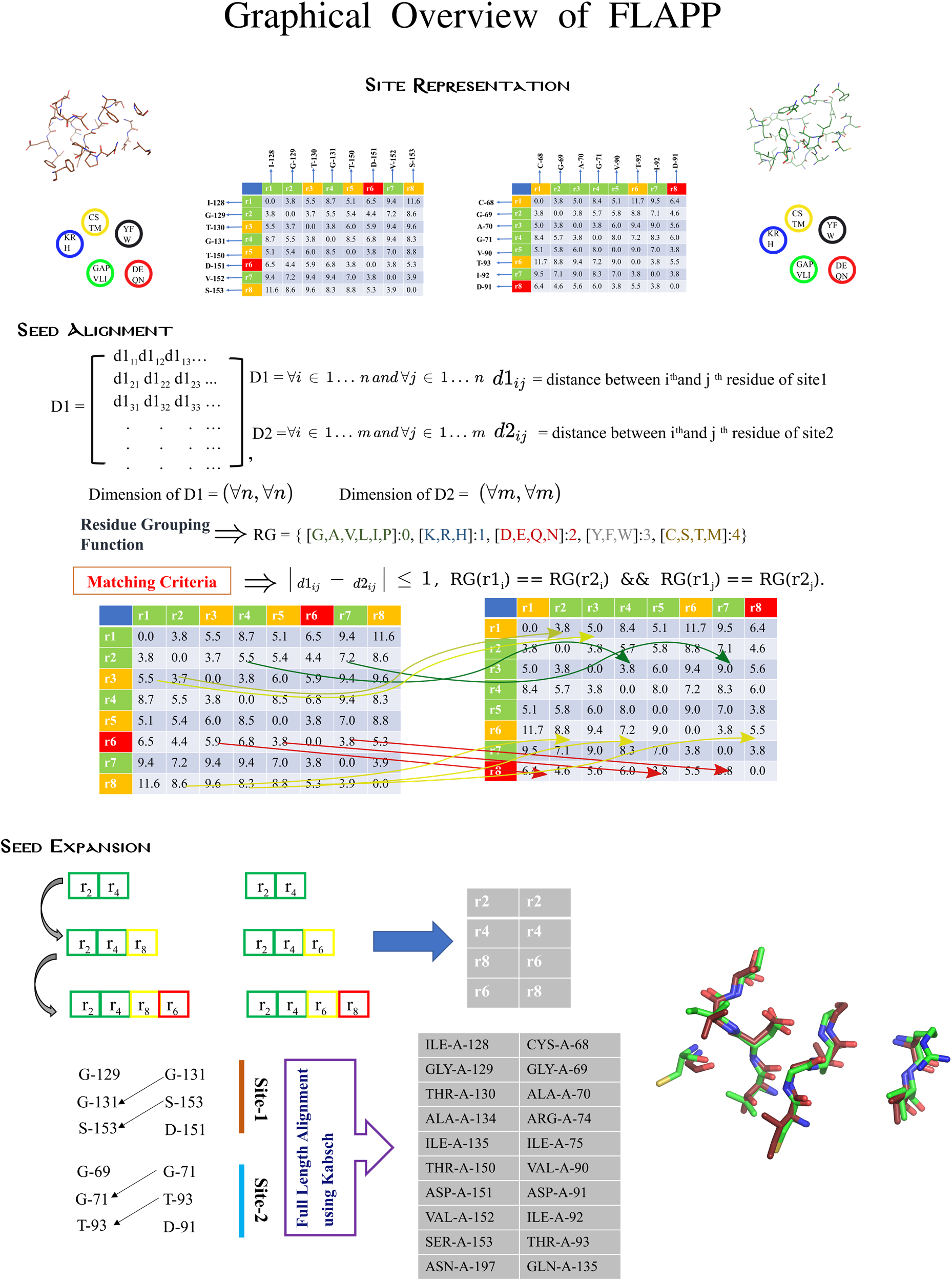
Graphical overview of the FLAPP algorithm. The FLAPP algorithm consists of three main modules: i) Binding site representation ii) Seed selection and iii) Seed expansion. FLAPP first represents binding sites as Cα distance matrices. Using physicochemical grouping and a distance metric from Eq. 1, it first selects residue pairs that are similar from both binding sites as seed primers. These primers are then extended with residues as long as the similarity criteria (i.e Eq. 1 and Eq. 2) is maintained. The seeds identified after the seed expansion step are used to get an optimal site rotation matrix via least square superposition using the Kabsch algorithm.

#### Site representation

The site representation module constructs a distance matrix for the binding site. For a given binding site of N residues, FLAPP first extracts Cα positions and constructs a distance matrix (D) of NxN dimensions that contains the euclidean distances for all N^2^ residue-pairs. Next, it groups all 20 amino acids into 5 standard groups based on the physicochemical properties. Group-0: (G,A,V,L,I,P); Group-1:(K,R,H); Group-2:(D,E,Q,N); Group-3(Y,F,W); Group-4 (C,S,T,M). The matrix D can also be thought of as a graph G(V,E), V representing the set of Cα atoms in each of the N residues and E representing all-pair distances among them.

#### Seed Primer selection

The second module exhaustively samples the sites to identify all possible ‘seeds’, which are progressively grown to find matching cliques. These are then screened to find the optimal alignment in the next module. From the adjacency matrices D1 and D2, it selects all distances that are similar based on two criteria: i) The magnitude of the difference between two distances is less than 1Å (Eq. 1), ii) The residues being compared belong to the same residue grouping (Eq. 2). Distances that satisfy these criteria are chosen as ‘seed primers’. The seed primers represent the common elements in a given pair of sites and by definition would form cliques.

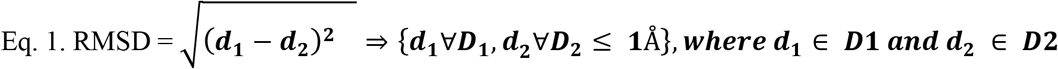

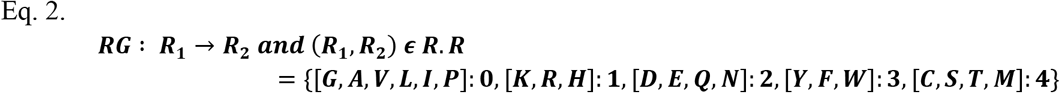

#### Seed Growth

At this point, each seed-primer contains two residues from each site based on Eq. 1 and Eq. 2. To add the next residue, we incorporate a strategy where we seek residues starting from the residue closest to the seed-primer. To accomplish this, we construct a sorted distance list for the trailing seed residue (D^sort^). Each row of D1^sort^ contains distances between any i^th^ residue of pocket-1 with all other residues in pocket-1. The program scans this list in the ascending order to find matching distances in D2^sort^. At this step, the seeds are progressively expanded as follows: an empty ArrayList is created to store the seeds and subsequent residues pairs are then added to the ‘ArrayList’ such that (a) the distances match according to our criteria (Eq. 1 and Eq. 2), (b) the newly added residues in both the structures have equivalent groups and (c) the residues in the ArrayLists from both the structures form matching cliques of size > 3. Residue pairs not associated with any clique are stored separately in a look-up table called the ‘HashTable’, which is consulted at each iteration to avoid repeated traversals of the same paths. The loop is iterated recursively until all residues are scanned. This step yields ‘expanded seeds’, which provide candidate alignments to be screened in the next step.

### 2.2 Least Square superposition using Kabsch algorithm

The candidate alignments are evaluated using the Kabsch least square superposition algorithm, from which the largest candidate (max n) with the least RMSD (<1Å) is taken as the optimal alignment of the two pockets. The Kabsch alignment is performed iteratively over all possible local alignments and only reports the maximum alignment^13^.

### 2.3 Alignment scores

The number of possible residues that can align between two binding sites is bounded by the size of the larger of the two sites being compared. In order to make comparison of binding site pairs more uniform, we define two scores: F_min_ and F_max_. F_min_ is defined as the ratio of the number of aligned residues to the size of the smaller binding site in the pair while F_max_ is defined as the ratio of the number of aligned residues to the size of the larger binding site in the pair.

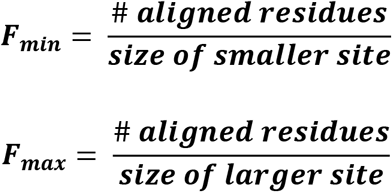

F_min_ and F_max_ are always expected to range between 0 and 1, both inclusive.

### 2.4 Code optimization

The algorithm was implemented using Python (Anaconda distribution 3.9). Many arithmetic operations such as calculating RMSD, Singular value decomposition (SVD) for Kabsch are implemented using numpy functions. For numerical calculations, Intel’s SVML (short vector math library) library has been utilized. The entire program was wrapped with Numba Just-In-Time compiler, which allowed Python code to be compiled into machine code specific to each system architecture (LLVM compiler)**^14^**. Moreover the code has been written in a format that facilitates Numba to compile into Single Instruction Multiple Data (SIMD) form. As a consequence of using Numba, the code had to be written such that the memory was pre-allocated for a few operations. For example, the ‘append’ function of python which adds new elements to a list is not supported by Numba. This is because the function ‘append’ creates a dynamic array whose size is not fixed and can be modified during execution. Incorporating such functions is not supported in compilation mode as we are required to provide the array size in advance. To circumvent this, a matrix of dimension 1000 was created with zero padding. The index of the array is then accessed and replaced sequentially via an incremental counter. The HashTable variable which is used to check if the cliques have been traversed before implements this logic.

The operations that further helped in optimizing the algorithm are: (i) the use of sorting at each seed selection which allows for considering local alignments as nearby residues are aligned first, (ii) creation of a LookUp table (size of n*100, where n represents the number of residues of site-1 and 100 corresponds to the index of the matched residues of site-2; At the start of execution, the LookUp matrix is initialized with zeros. For every match detected, a counter variable was incremented to one which will replace zeros with the index pointing to the residue number of site-2) to keep track of paths that have already been explored and hence avoid futile traversals, (iii) use of the Intel SVML library for computing the dot products in the construction of rotation matrices for the calculation of RMSD.

## Results

We present a fast and accurate algorithm *FLAPP* for aligning binding sites in protein structures. An alignment of a pocket-pair (25 residues each) takes 12 ms (~1/80 s) on a single desktop processor (Intel i5-8400 2.80GHz). The speed of obtaining alignments at the atomic level enables accurate searching at a PDB-scale.

### 3.1.1 Robustness of FLAPP to minor perturbations in binding site

To evaluate the robustness of FLAPP to geometric and compositional changes in binding sites, we systematically tested the (i) sensitivity of FLAPP to perturbations in binding site residue positions and (ii) sensitivity of FLAPP to mutations in binding site residues. In order to ensure that our analysis is not biased by the choice of ligand, we systematically selected a diverse set of sites (corresponding to 19 ligands) by constructing a 2D matrix based on molecular weights and partition coefficients of all ligands in PDB and selecting representatives from different regions (Fig 2a).

**Fig. 2.**
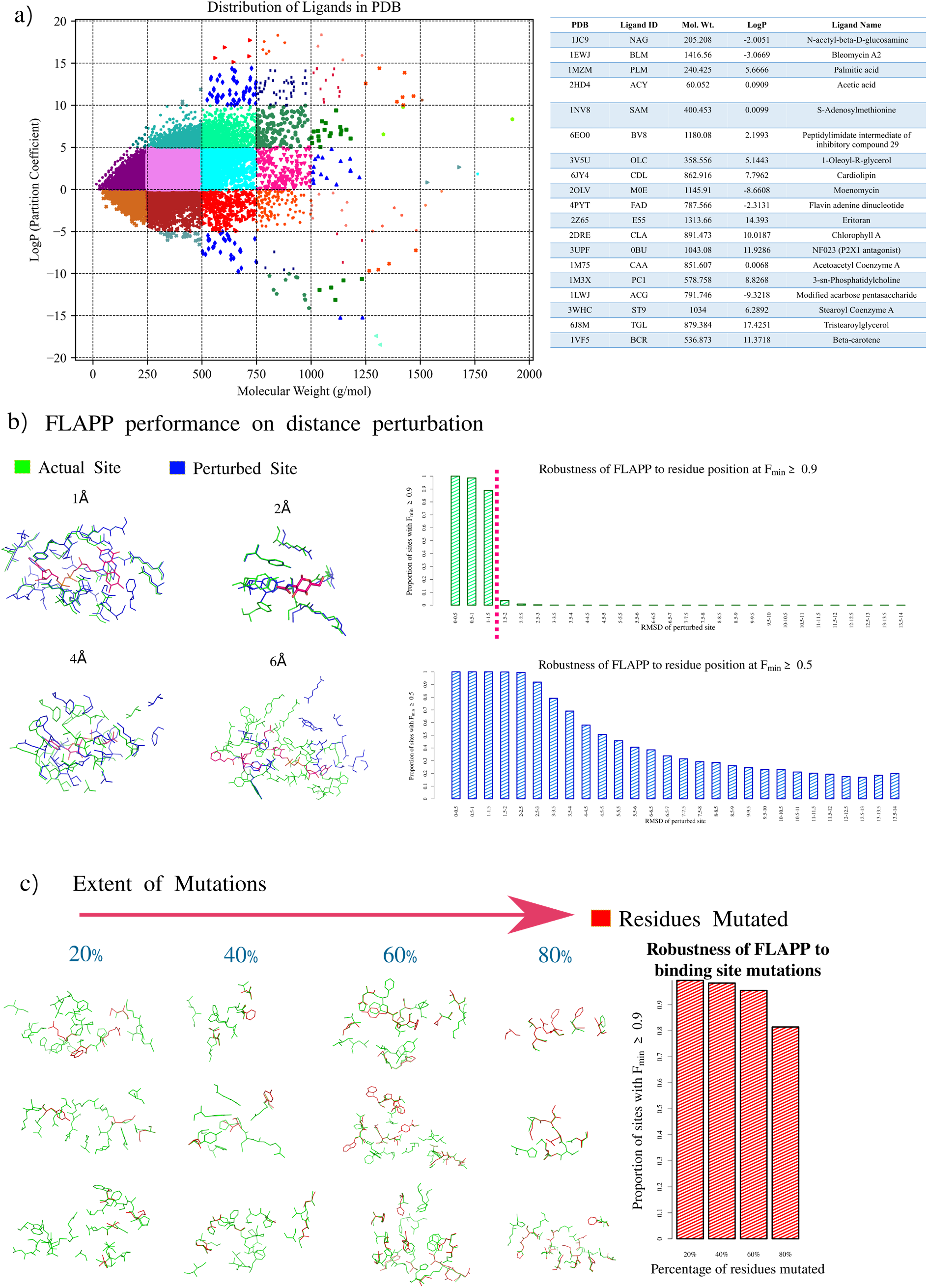
Assessing the sensitivity of FLAPP to different perturbations. a) Grouping of ligands in PDB based on molecular weight (g/mol) and partition coefficient (LogP). The plot is partitioned into grids of 250 g/mol on the x-axis and 5 units of LogP on the y-axis. 19 diverse ligands were chosen and one binding site for each was chosen randomly. b) The residues in the 19 binding sites were randomly perturbed so as to generate 3000 synthetic sites each, with RMSDs ranging from 0 - 14Å. The sensitivity of FLAPP in aligning residues in the original site with the perturbed sites is explored at two F_min_ scores (0.5 and 0.9). c) Similarly, synthetic binding sites that had a proportion of their residues mutated were generated for each of the 19 binding sites.

We first tested FLAPP’s tolerance to perturbations by synthetically generating 3000 new binding sites from each of the 19 diverse sites by randomly perturbing their residue positions. The perturbations were done such that the newly generated binding sites had a bounded RMSD ranging between 0 to 14Å with respect to the original site. Site alignments were then evaluated by FLAPP for each perturbation. FLAPP aligns perturbed sites with their parent sites with perfect F_min_ and F_max_ scores up to an RMSD of 1.5Å in all cases (Fig 2b) indicating that it is robust to minor perturbations in residue positions. Our analysis mimics perturbations seen in real life scenarios that can occur due to minor crystallographic variations or conformational changes due to ligand binding or even positional flexibility in NMR data. As the perturbations cross 6Å, the scores drop below 0.5. This is expected as the RMSD increases, residue-to-residue alignments become weaker.

Next, we tested the sensitivity of FLAPP to change in binding site residue types, by systematically generating mutations in the site. For each of the 19 binding sites, we generated 1000 new sites for each site by randomly mutating some proportion of residues to other residues. FLAPP successfully aligns sites even after mutating up to 60% of their residues (Fig 2c). These perturbations are designed to mimic comparison of sites in homologous proteins and situations such as polymorphisms and disease associated mutations, which are typically seen only in a small portion of binding sites. The fact that our algorithm is robust upto 60% of changes in the site, implies that residue grouping does not adversely affect its performance in achieving optimal alignments. The use of the Kabsch algorithm after selection of the seeds ensures that residue-residue correspondences missed due to grouping are salvaged at the final alignment step (Supplementary File: PerturbationAnalysis-Supplementary.xlsx).

### 3.1.2 Evaluating FLAPP’s accuracy across levels of increasing complexity of comparison

In order to systematically evaluate the alignment accuracy, FLAPP was tested on diverse pairs of binding sites. Here we grouped binding sites into three levels of complexity (Easy, Medium and Hard targets) based on sequence and structure similarities. (A) The easy targets include 7563 protein pairs from pdb_95 that share high sequence identity (>95%), belong to the same SCOP family and bind to the same ligand^15^. As the proteins are nearly identical, their binding sites are expected to be the same. FLAPP successfully showed identical alignments in 98% (F_min_ ≥ 0.8 and alignment length > 10) (Supplementary Fig 1a and Supplementary File: Level-Supplementary.xlsx). (B) the medium complexity targets were built using 729 protein pairs which do not have any similarity in sequence (identity <30%), but exhibit structural similarity (belong to the same SCOP superfamily)^16^. The idea is that structurally-similar and sequentially-dissimilar proteins binding to the same ligand tend to share commonality in their binding sites. Out of 729, 416 pairs consist of binding sites that bind to the same ligand, which we consider as true positives. The remaining 313 are binding sites recognizing different ligands (Tanimoto coefficient < 0.5) and are considered to be negative controls. FLAPP achieves an ROC of 0.93 in this dataset, indicating that it can successfully handle binding sites from structurally similar, but sequentially dissimilar proteins (Supplementary Fig 1b and Supplementary File: Level-Supplementary.xlsx). (C) The hard targets dataset contains protein pairs that are diverged both at sequence and the structure but recognise common ligands. Because it is the same ligand that is getting recognised, the proteins could possess some commonality in their site. We took such proteins from previously published literature (Ausiello *et al*.) where the authors explored a case of convergent evolution and reported 4 pairs of proteins that are structurally distinct but possess good commonality at their binding sites^17^. The mentioned pairs also intriguingly display sequence order reversal at the binding site which conventional sequence and structural alignment approaches fail to align. FLAPP is able to successfully align all these pairs. The residue-residue correspondence reported by our method is the same as that described by the authors. The optimal alignments reported by FLAPP for these pairs are shown in Fig 3. This in fact illustrates the importance of site based alignment over traditional sequence or structural comparison. In addition to these targets at three levels, we also constructed a dataset of 50,000 binding site pairs that are dissimilar (sequence identity < 30%, different SCOP folds) and bind to different ligands (Tanimoto Coefficient < 0.5). These serve as negative controls for evaluating specificity. FLAPP did not yield any significant alignment amongst these pairs, indicating high specificity (Supplementary File: Level-Supplementary.xlsx). Put together, our results show that FLAPP has high sensitivity and specificity.

**Fig. 3.**
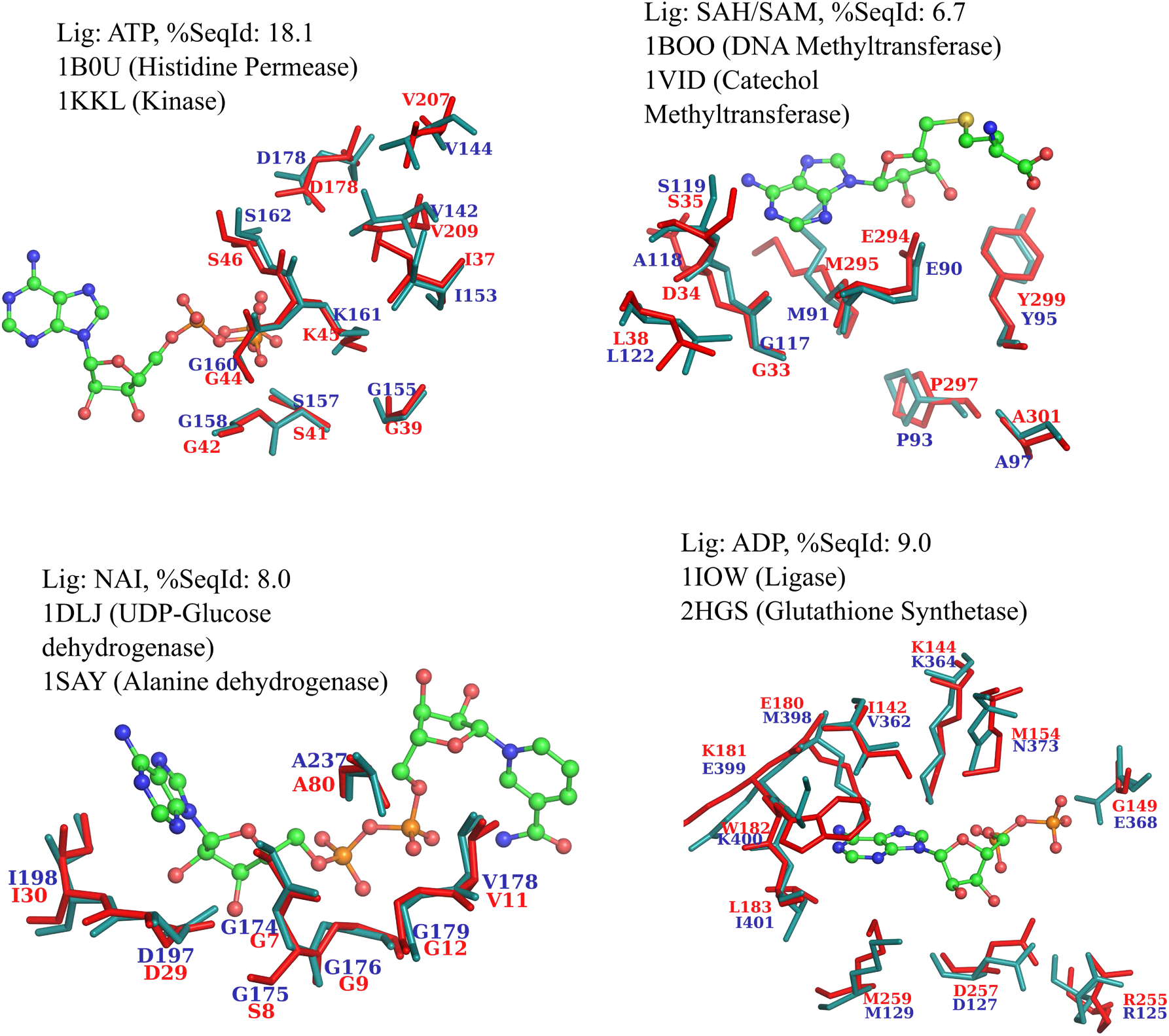
Alignments produced by FLAPP in the hard targets validation set. Each binding site pair has less than 20% sequence identity and proteins belong to different folds. All 4 pairs have sequence order reversal in the sequence. FLAPP successfully identifies similarities within each pair and outputs accurate residue-residue matches.

### 3.1.3 Benchmarking of accuracy and sensitivity of FLAPP against other methods

Next, we benchmark the accuracy of FLAPP against two site alignment algorithms, G-LoSA and SiteMotif^7,8^, that are currently the best alignment-based algorithms. Other methods that produce optimal alignments, eg., PocketAlign and SiteEngine, are not included as they need around five minutes per comparison. G-LoSA and SiteMotif on average take less than a second. To enable unbiased, systematic analysis, we assess the alignment accuracy of FLAPP, SiteMotif and G-LoSA on two datasets of non-redundant binding sites. Dataset-1 comprises a set of 514 pairs of proteins that bind to the same ligand and belong to the same SCOP superfamily but different SCOP families. This is essentially an obvious case since the sequence as well as the structures share a high degree of similarity at the SCOP superfamily level. As expected all three methods fare very well on this dataset (Fig 4a). Dataset-2 comprises a pair of proteins that binds to similar ligands but belong to different SCOP folds. This dataset can be considered as inherently difficult as the proteins share no similarity in the sequence and the structure. We compute the length of alignment reported by each method as our metric of accuracy. SiteMotif shows the best performance in Dataset-2, followed closely by FLAPP and then G-LoSA (Fig 4b). However, FLAPP and SiteMotif consistently achieve longer alignments compared to G-LoSA(Supplementary File: Datasets-Supplementary.xlsx.)

**Fig. 4.**
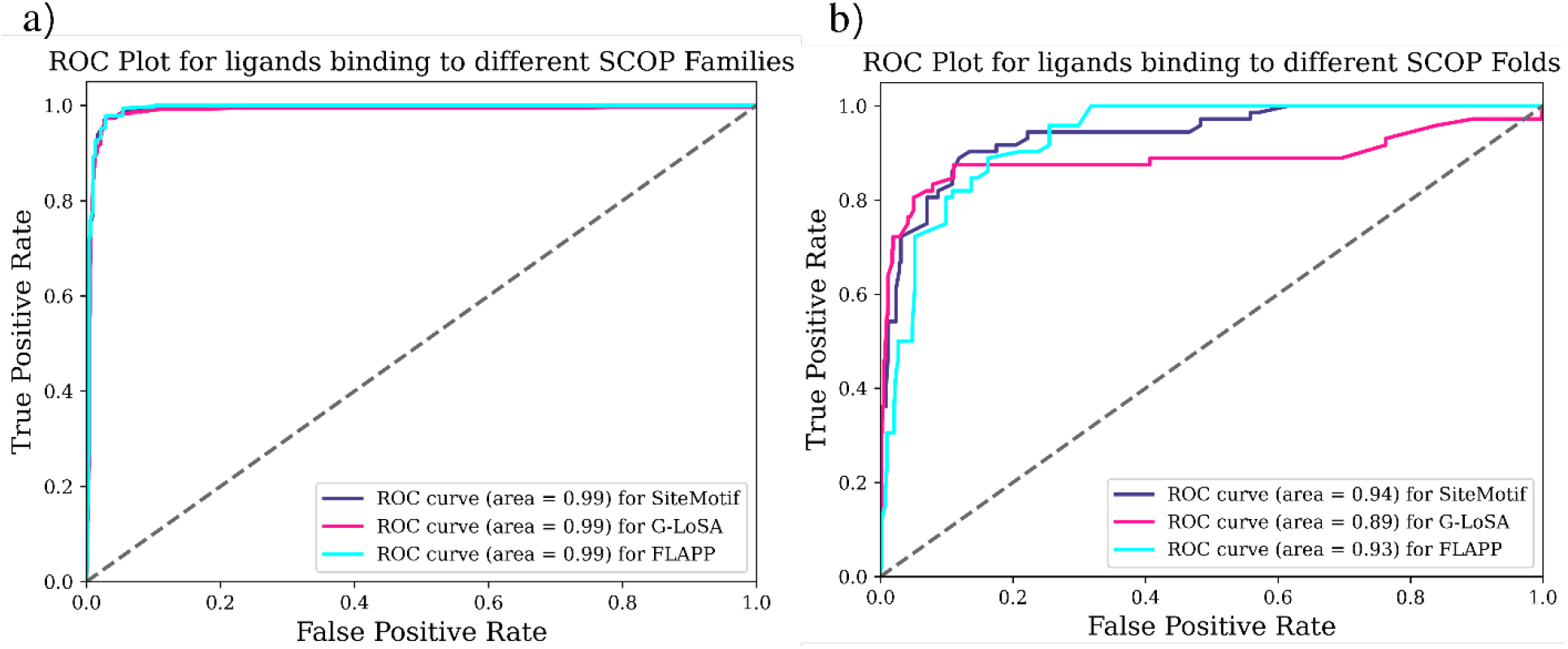
Alignment accuracy of FLAPP over known site alignment tools (SiteMotif and G-LoSA). Two sets of binding site pairs are analyzed, a) 2421 pairs, of which 514 are true positives (similar sites) and 1907 are true negatives. All three methods are seen to fare well; b) binding sites that exhibit only partial similarity between them capturing the length of alignment of each method, 5383 pairs are considered of which 72 are true positives and 5311 true negatives.

### 3.2 Runtime Performance

#### 3.2.1 Benchmarking the runtime of FLAPP

Traditional alignment based site comparison methods, despite being accurate, are often many orders of magnitude slower than alignment free methods. This is because alignment operations such as rotation, translation and least squares superpositions are computationally expensive. The FLAPP algorithm and implementation is designed to mitigate these factors. Large scale analyses of binding sites typically involve the use of alignment-free methods such as PocketMatch and RAPMAD because of their quick execution times of ~ 1/250s and ~1/100s for comparing a site pair. A comparison of 1 million binding site pairs is expected to complete in around 70 minutes. In comparison, SiteMotif or G-LoSA, would take around 6 days. Alignment-free methods output a score that represents the similarities between the sites while alignment-based methods provide us with full length residue-residue correspondences. FLAPP bridges the gap between these sets of tools, bringing in the accuracy of alignment-based methods while competing with the speed of alignment-free methods. To obtain an accurate estimation of the performance, FLAPP was measured against different sets ranging from 10^2^ to 10^6^ sites taken randomly from PDB. The performance was timed with G-LoSA and SiteMotif. G-LoSA can be executed in one of two ways: either with or without file pre-processing. We term the former as ‘G-LoSA-Preprocessed’ and the latter as ‘G-LoSA-Native’. The native version processes the input PDB files on each run using a Java program while the input files are processed beforehand in the preprocessed version. From Fig 5a, it is clear that FLAPP tops the list as the fastest alignment program, outperforming both SiteMotif and G-LoSA by a huge margin. Even the preprocessed version of G-LoSA is 4x slower than our method.

**Fig 5.**
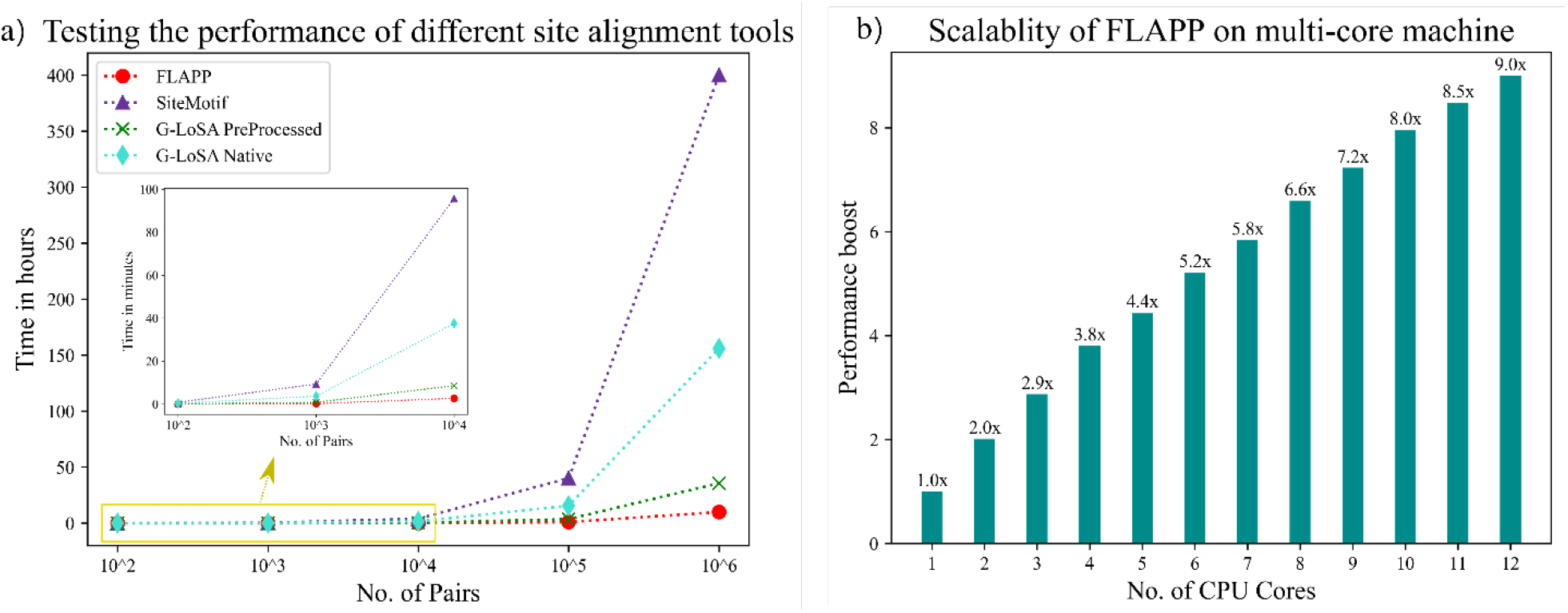
Execution time of FLAPP over known site alignment. a) Performance comparison of FLAPP, SiteMotif and G-LoSA on different dataset sizes. The runtimes of FLAPP, G-LoSA (Preprocessed and Native) and SiteMotif were compared on sets of randomly selected binding site pairs. Sets of different sizes varying from 100 pairs to 0.1 million pairs were used to test the scalability of the method. FLAPP was able to align 1,000,000 pairs in 1.9 hours (~ 6ms per alignment). Our method is able to achieve at least 4X performance compared to the closest competitor G-LoSA PreProcessed and 20X faster than the actual G-LoSA program while maintaining accuracy on par with the SiteMotif. b) Scalability of FLAPP to multiple CPU cores: The multi-processing enabled version of FLAPP was run on many cores and the performance boost was obtained. FLAPP efficiently utilizes all available CPU cores in order to scale further. At 12 cores, FLAPP aligns 1080 pairs per second (~ 1ms per pair).

#### 3.2.2 A parallelized implementation scales well across multiple CPU cores

The speed benchmarking analysis showed that FLAPP on average takes 10 milliseconds to align a pair of sites on a single desktop computer. In other words, the single threaded version of FLAPP aligns 120 pairs per second. This is by far the fastest implementation of alignment-based site comparison softwares that exists. However, modern CPUs are designed to have many cores which permits concurrent execution of individual statements across multiple processes. We therefore aimed to utilize all available CPUs to further accelerate our algorithm. FLAPP code is wrapped in a multiprocessing module that will assign each pair of pockets to a different computing resource. To assess the performance, we randomly generated 1 million pairs of binding sites from the PDB for which serial implementation took 140 minutes to complete. The performance of the parallel implementation was evaluated as the ratio of the time taken by the serial version (T_s_) to time taken by parallel version (T_p_) at different no. of CPU cores (N), where N ≥ 1 (Fig 5b). At N=2, FLAPP runs approximately 2x faster than the serial version. This implied that FLAPP can scale with an increasing number of CPUs. Scaling was however not linear since the multiprocessing introduces CPU overheads. Utilizing 12 cores, FLAPP was able to align 1080 site pairs per second, a ninefold improvement over the serial version. At N=12, FLAPP sets a new benchmark as the first alignment-based approach that needs only 1 millisecond to compare two sites. A standard desktop machine equipped with a Intel Core i7-10700 was used for this analysis.

### 3.3 New application capabilities that emerge out of the speed

#### Case Study: Analysis of Vitamin B12 binding sites

We showcase the performance of FLAPP in achieving a PDB-wide scan of 3D binding sites for Vitamin B12 binding sites as an example. Vitamin B12 is a large ligand and is known to bind to diverse protein structures^18^. We performed a two level systematic analysis: i) To find a Vitamin B12 binding motif and (ii) To scan the motif against PDB. First we SiteMotif, a recently developed method from our group for multiple site alignment. A total of 83 proteins were extracted from the PDB that are complexed with Vitamin B12. Identical sequences within them were removed using a sequence identity cut off 95 which resulted in 51 proteins. We then removed redundancies at structural levels by doing all-to-all 3D structural alignment using the TM-align program^19^. TM-align, in addition to doing structural alignment, returns a TM-Score to numerically quantify the extent of similarity between proteins. As reported by the authors, TM-Score > 0.5 indicates that structures being compared are likely to be similar. We used the same cutoff and grouped all 51 proteins into 12 structure-based clusters, which indicated that the Vitamin B12 ligand binds to proteins from 12 different structural families (Supplementary File B12-Supplementary.xlsx). Next, we checked if the sites of 12 proteins share any commonality in their binding sites using FLAPP. We observed that 7 of the 12 share significant similarity in their sites (alignment length > 10, RMSD <1Å) (Supplementary Fig 2), despite belonging to different SCOP folds, suggesting this to be a predominant binding site type for binding Vitamin B12. We examined the binding sites for the remaining 5 clusters and found that they were very different from the other groups, and as they formed singletons, we did not consider them for further analysis. The selected 7 Vitamin B12 proteins are 1XRS, 2REQ, 3IV9, 3KOX, 5C8A, 5CJV and 6WTE, showed an average sequence identity of 20.8% among them and a mean structural deviation of 18.2Å, in consonance with their diversity. We then examined the positional conservation among these using SiteMotif (by aligning each of them onto 3KOX taken as a representative). This yielded a structural motif (Fig 6) that contained a histidine coordinating with the cobalt, an aspartic acid and serine. These residue conservation with Vitamin B12 has been reported earlier^18,20^.

**Fig 6.**
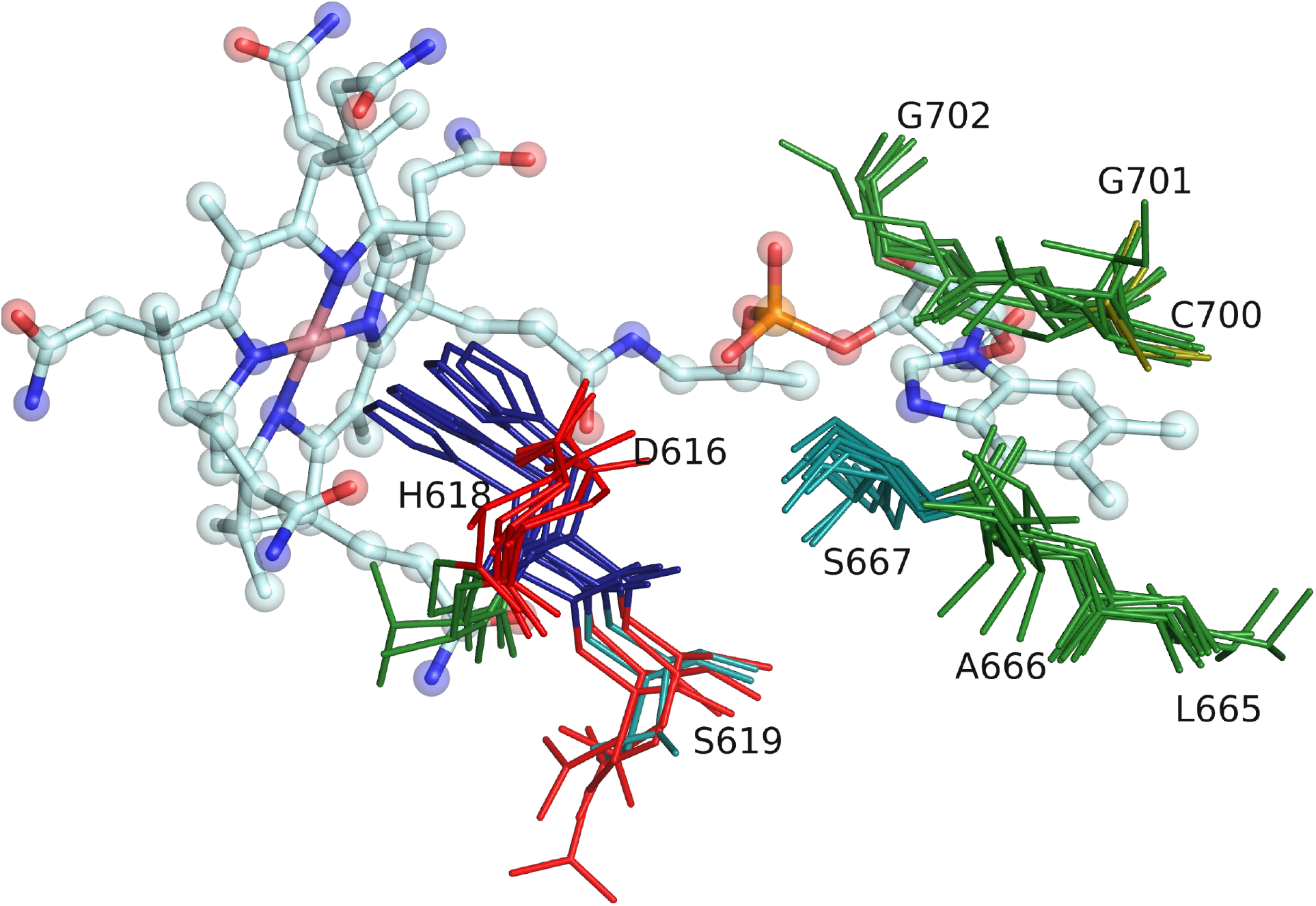
Deriving the structural motif for Vitamin B12 binding sites. For the set of 7 unique Vitamin B12 proteins, the binding sites are extracted and the conserved residues are identified using SiteMotif. The motif thus derived is used with FLAPP for large scale analysis such as a PDB-wide search. Vitamin B12 is represented in ball and stick and the conserved residues are represented in sticks. Residues are colored according to their physicochemical property.

We then performed a one-vs-all binding site alignment between the Vitamin B12 motif and the entire set of known binding sites from PDB using FLAPP. FLAPP successfully recovered all 7 known binding sites containing the motif. We tested if the motif is present in all Vitamin B12 binding proteins including those in homologous proteins that are not crystallized with Vitamin B12. To do this, we established an unbiased scan of the Vitamin B12 structural motif against pockets from an extensively augmented pocketome PocketDB that we have previously developed^21^. PocketDB, consists of high confidence putative pockets for all structures in PDB (PDBv-2014) using pocket prediction algorithms (1,39,718 pockets). FLAPP scanned the Vitamin B12 motif against 139,718 pockets of PocketDB (9,78,026 alignment operation). FLAPP completed the scan on a 12 core machine in 10 minutes. FLAPP reported 26 hits for the Vitamin B12 motif. These 26 proteins were known to utilize Vitamin B12 for its function but the structures complexed with Vitamin B12 are available only for 12 of them. This indicates that FLAPP was able to correctly identify the apo form of the Vitamin B12 binding proteins. An exhaustive site alignment exercise, such as one which involves PDB level scale, will provide a number of insights that were previously considered to be impossible due to the inherent complexity of the current algorithms during runtime. Here we demonstrate that FLAPP can perform such analysis on a normal desktop system in reasonable time.

## Discussion

Biological reactions are driven by biomolecular interactions. Elucidating the characteristics of interactions governing between proteins and ligands can provide functional insights as to why cells behave the way they do or which proteins a particular ligand can bind to. The demand to address such questions has gained significant attention as it drives us to demystify novel protein targets and cellular pathways that are currently unknown to science. This is even more important if the ligand being investigated is a drug molecule or a probable lead candidate. Often, precise identification of small molecule binding sites provide direct answers into understanding the mechanism by which proteins recognize ligands.

We present here a highly scalable algorithm for binding site comparison that provides fast and accurate residue-level alignments and facilitates rapid identification of similar binding sites at a proteome level. Since binding sites provide a direct window into understanding the function of proteins, this opens up the possibility for extensive protein annotation by knowledge transfer based on site similarity. Alignments at binding site level allow us to probe similarity at residue level, providing us with much higher confidence in our annotation process. PDB currently (v 30 April 2022), contains 189,915 structures - of which 68,968 are crystallized with specific ligands^15^. Furthermore, pocket prediction methods will expand the repertoire of potential ligand recognition sites by several fold^10,11,21^.

FLAPP was tailor made for binding site comparisons at a scale that meets the pace at which data is being generated. Our benchmarks show that FLAPP produces alignments comparable to the existing state-of-the-art at blazing speed of 1/80th of a second per comparison. FLAPP accomplishes this speed as a consequence of various optimizations at the implementation level which includes, grouping of residues, hash table to keep track of traversed path, SIMD instructions whenever possible, and lastly the use of compiler libraries to compile the whole program based on system specific architecture. In addition, FLAPP supports multiprocessing and can harness the power of multiple CPU cores to provide alignments at a speed of about one millisecond per pair on a standard 12 core desktop computer.

Similarities at the binding site level can be used in both protein-centric fashion to identify potential ligands that can bind to it and in a ligand-centric fashion to identify different proteins that can potentially bind the small molecules. The most obvious use of such a capability is to annotate ligand-binding ability at a proteome-wide scale. In addition, binding site alignments have the potential for immense application in various aspects of drug discovery - specifically in drug repurposing and in identifying drug off-targets.

## Supporting information

PerturbationAnalysis-Supplementary

Level-Supplementary

## Acknowledgement

We acknowledge support from the Bioinformatics grant, Department of Biotechnology, Government of India. Naren Chandran Sakthivel is funded by the Council of Scientific & Industrial Research, Government of India (Senior Research Fellow - File no: SPM-07/079(0287)/2019-EMR-I).

## Author Contributions

Santhosh Sankar (SS) and Naren Chandran Sakthivel (NCS) developed and validated the algorithm under the supervision of Nagasuma Chandra (NC). SS implemented the algorithm. SS and NCS carried out the analysis. All authors have read and approved the manuscript.

## Supplementary Figures

**Supplementary Fig 1.**
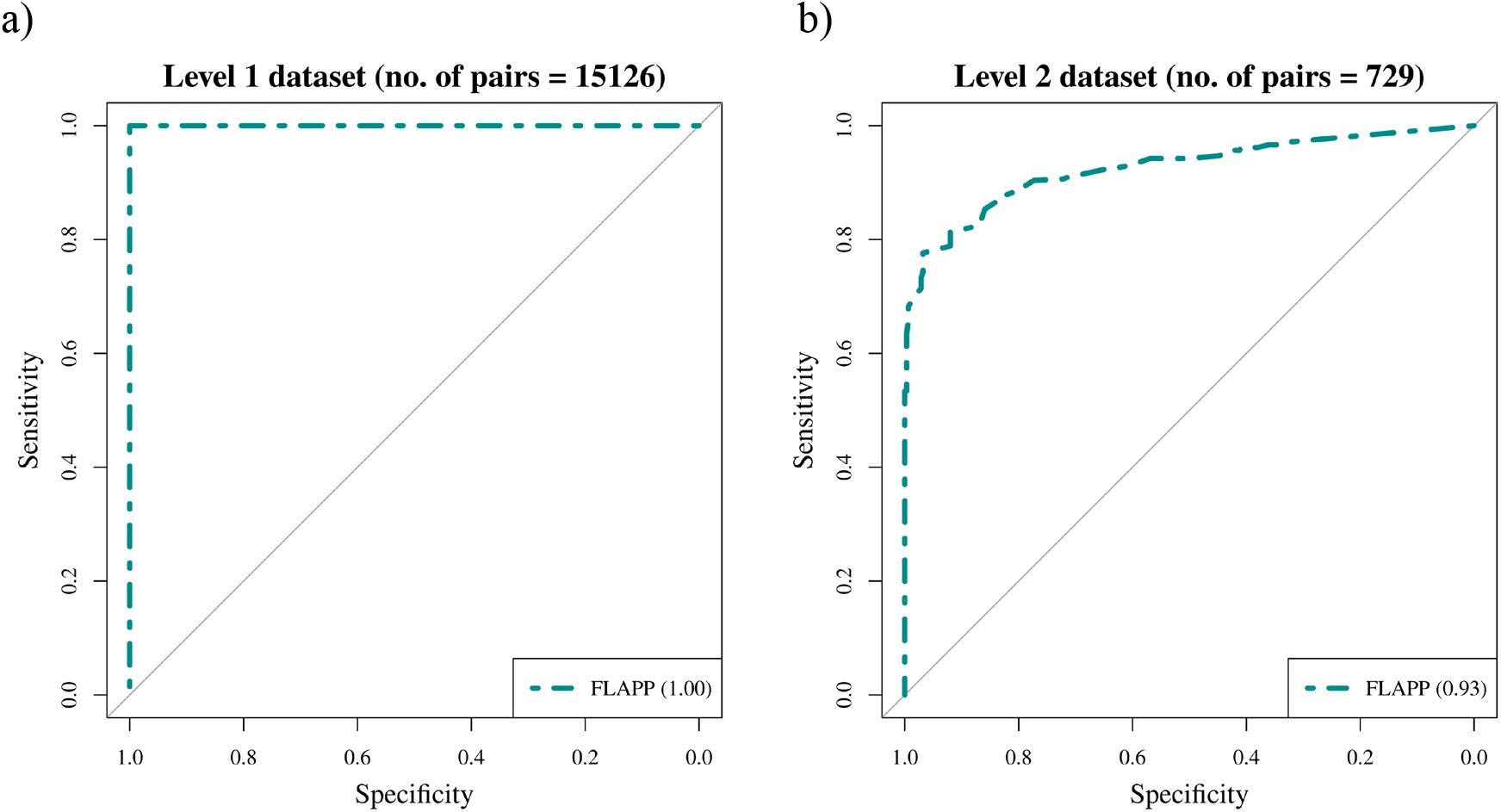
The accuracy of FLAPP was assessed using datasets constructed to query the ability to find alignment in specific scenarios. (a) Positive binding site pairs were constructed by pairing proteins from PDB that have >95% sequence identity and bind to the same ligand. Negative controls were constructed from the positive binding pairs by perturbing residue positions such that the overall RMSD for each site was greater than 6Å. This is a trivial case and as expected, FLAPP achieves perfect accuracy. (b) Positive binding site pairs were constructed by pairing proteins from PDB that belong to different SCOP families, but same SCOP superfamily but bind to the same ligand. Negative binding site pairs were selected using the same kind of SCOP differences, but with the criteria that they did not bind to the same ligand.

**Supplementary Fig 2.**
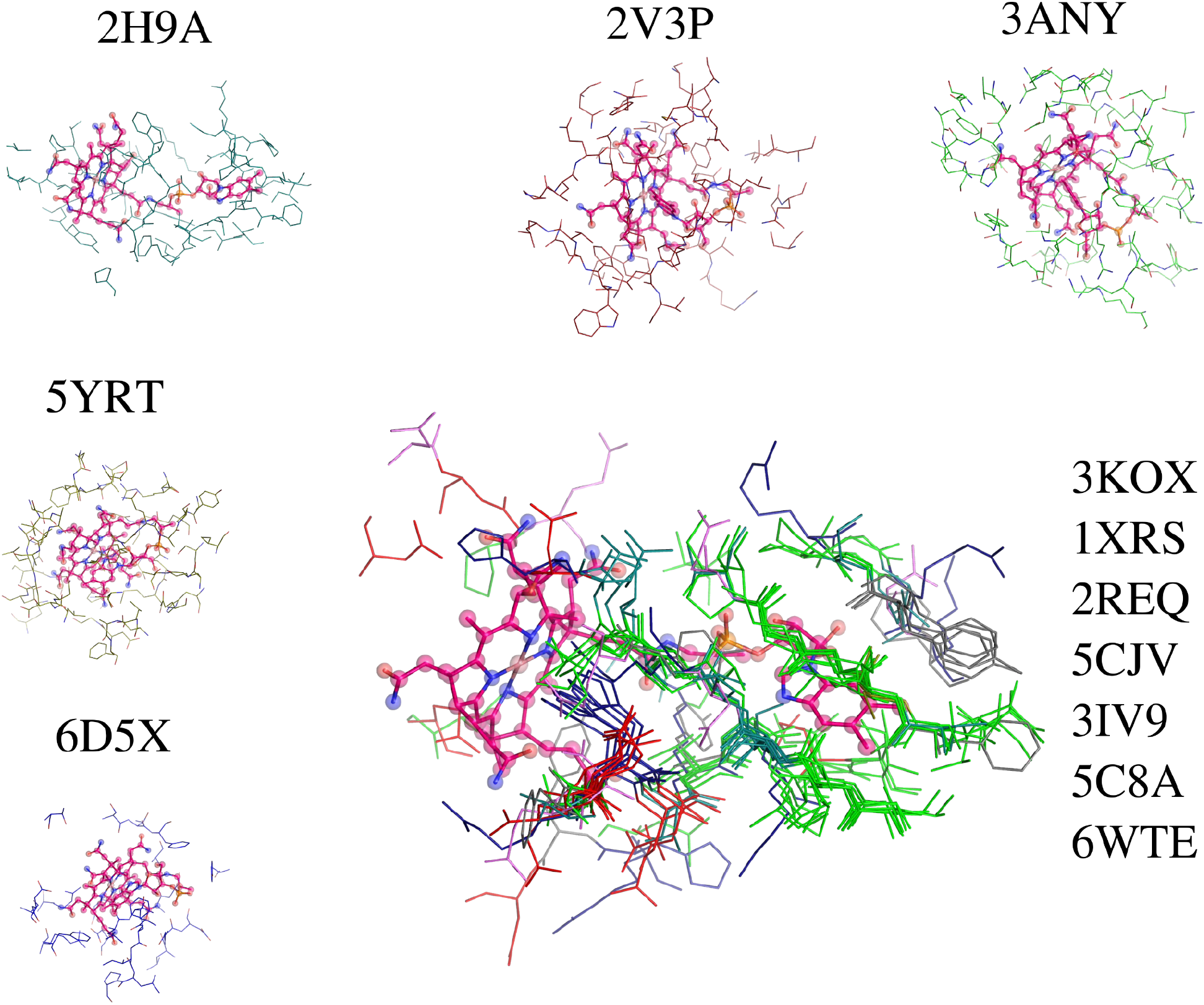
Binding site similarity of the 12 selected Vitamin B12 binding site representatives. Showing the multiple site alignment of superposing 7 sites onto representatives (3KOX) using SiteMotif. The 7 representatives that shared similarity are shown as aligned structures. The remaining 5 representatives are also shown separately. Vitamin B12 ligand is represented in a ball and sticks in pink.

## Notes

https://github.com/santhoshgits/FLAPP

### Competing Interest Statement

The authors have declared no competing interest.

## References

1. Loewenstein, Y. et al. Protein function annotation by homology-based inference. Genome Biol. 10, 207 (2009).

2. Altschul, S. F., Gish, W., Miller, W., Myers, E. W. & Lipman, D. J. Basic local alignment search tool. J. Mol. Biol. 215, 403–410 (1990).

3. Yeturu, K. & Chandra, N. PocketMatch: A new algorithm to compare binding sites in protein structures. BMC Bioinformatics 9, 543 (2008).

4. Krotzky, T., Grunwald, C., Egerland, U. & Klebe, G. Large-Scale Mining for Similar Protein Binding Pockets: With RAPMAD Retrieval on the Fly Becomes Real. J. Chem. Inf. Model. 55, 165–179 (2015).

5. Desaphy, J., Raimbaud, E., Ducrot, P. & Rognan, D. Encoding Protein–Ligand Interaction Patterns in Fingerprints and Graphs. J. Chem. Inf. Model. 53, 623–637 (2013).

6. Yeturu, K. & Chandra, N. PocketAlign A Novel Algorithm for Aligning Binding Sites in Protein Structures. J. Chem. Inf. Model. 51, 1725–1736 (2011).

7. Lee, H. S. & Im, W. G-LoSA: An efficient computational tool for local structure-centric biological studies and drug design: G-LoSA. Protein Sci. 25, 865–876 (2016).

8. Sankar, S. & Chandra, N. SiteMotif: A graph-based algorithm for deriving structural motifs in Protein Ligand binding sites. PLOS Comput. Biol. 18, e1009901 (2022).

9. Shulman-Peleg, A., Nussinov, R. & Wolfson, H. J. SiteEngines: recognition and comparison of binding sites and protein-protein interfaces. Nucleic Acids Res. 33, W337–W341 (2005).

10. Kalidas, Y. & Chandra, N. PocketDepth: A new depth based algorithm for identification of ligand binding sites in proteins. J. Struct. Biol. 161, 31–42 (2008).

11. Le Guilloux, V., Schmidtke, P. & Tuffery, P. Fpocket: An open source platform for ligand pocket detection. BMC Bioinformatics 10, 168 (2009).

12. Ghersi, D. & Sanchez, R. Improving accuracy and efficiency of blind protein-ligand docking by focusing on predicted binding sites. Proteins Struct. Funct. Bioinforma. 74, 417–424 (2009).

13. Kabsch, W. A solution for the best rotation to relate two sets of vectors. Acta Crystallogr. Sect. A 32, 922–923 (1976).

14. Lam, S. K., Pitrou, A. & Seibert, S. Numba: a LLVM-based Python JIT compiler. in Proceedings of the Second Workshop on the LLVM Compiler Infrastructure in HPC - LLVM ’15 1–6 (ACM Press, 2015). doi:10.1145/2833157.2833162.

15. Berman, H. M. The Protein Data Bank. Nucleic Acids Res. 28, 235–242 (2000).

16. Lo Conte, L. SCOP: a Structural Classification of Proteins database. Nucleic Acids Res. 28, 257–259 (2000).

17. Ausiello, G., Peluso, D., Via, A. & Helmer-Citterich, M. Local comparison of protein structures highlights cases of convergent evolution in analogous functional sites. BMC Bioinformatics 8, S24 (2007).

18. Sukumar, N. Crystallographic studies on B12 binding proteins in eukaryotes and prokaryotes. Biochimie 95, 976–988 (2013).

19. Zhang, Y. TM-align: a protein structure alignment algorithm based on the TM-score. Nucleic Acids Res. 33, 2302–2309 (2005).

20. Tollinger, M., Konrat, R., Hilbert, B. H., Marsh, E. N. G. & Kräutler, B. How a protein prepares for B12 binding: structure and dynamics of the B12-binding subunit of glutamate mutase from Clostridium tetanomorphum. Structure 6, 1021–1033 (1998).

21. Bhagavat, R., Sankar, S., Srinivasan, N. & Chandra, N. An Augmented Pocketome: Detection and Analysis of Small-Molecule Binding Pockets in Proteins of Known 3D Structure. Structure 26, 499–512.e2 (2018).

